# Rapid cross-sensory adaptation of self-motion perception

**DOI:** 10.1101/2021.06.16.448688

**Authors:** Shir Shalom-Sperber, Aihua Chen, Adam Zaidel

## Abstract

Perceptual adaptation is often studied within a single sense. However, our experience of the world is naturally multisensory. Here, we investigated cross-sensory (visual-vestibular) adaptation of self-motion perception. It was previously found that relatively long visual self-motion stimuli (≳ 15s) are required to adapt subsequent vestibular perception, and that shorter duration stimuli do not elicit cross-sensory (visual↔vestibular) adaptation. However, it is not known whether several discrete short-duration stimuli may lead to cross-sensory adaptation (even when their sum, if presented together, would be too short to elicit cross-sensory adaptation). This would suggest that the brain monitors and adapts to supra-modal statistics of events in the environment. Here we investigated whether cross-sensory (visual↔vestibular) adaptation occurs after experiencing several short (1s) self-motion stimuli. Forty-five participants discriminated the headings of a series of self-motion stimuli. To expose adaptation effects, the trials were grouped in 140 batches, each comprising three ‘prior’ trials, with headings biased to the right or left, followed by a single unbiased ‘test’ trial. Right, and left-biased batches were interleaved pseudo-randomly. We found significant adaptation in both cross-sensory conditions (visual prior and vestibular test trials, and vice versa), as well as both unisensory conditions (when prior and test trials were of the same modality – either visual or vestibular). Fitting the data with a logistic regression model revealed that adaptation was elicited by the prior stimuli (not prior choices). These results suggest that the brain monitors supra-modal statistics of events in the environment, even for short-duration stimuli, leading to functional (supra-modal) adaptation of perception.

## INTRODUCTION

To act in the world, humans and animals need to maintain a veridical percept of their position, orientation, and movement, relative to the environment. Thus, accurate perception of one’s motion in space (self-motion perception) is a vital skill. However, the environment itself is dynamic, making this a challenging task for the brain. Moreover, repeated sensory stimuli do not elicit the same neuronal responses. Rather, these adapt, often leading to altered perception (aftereffects). For example, the motion aftereffect: exposure to a continuous visual motion stimulus in one direction leads to sensory adaptation, and a subsequent bias towards perceiving visual motion in the opposite direction (Anstis, Verstraten, & Mather, 1998; Thompson, P., & Burr, 2009). Similarly, the vestibular motion also leads to adaptive aftereffects (Crane, 2012; Gordon, Fletcher, Melvill Jones, & Block, 1995). Adaptation may come at the cost of perceptual accuracy. But, it also has benefits. Specifically, adaptation increases sensitivity to novel stimuli and improves efficiency by not wasting cognitive and energy resources on constant or uninformative signals (Wark, Lundstrom, & Fairhall, 2007). Either way, whether it is overall beneficial or not, adaptation continuously and substantially influences our perception and is thus an important aspect of brain research.

Sensory adaptation is ubiquitous, and has been described in many sensory modalities (Dalton, 2000; Di Lorenzo & Lemon, 2000; Kohn, 2007; Schweinberger et al., 2008). However, perceptual experiences are seldom uni-sensory. Rather, we generally interact with the world using multiple sensory modalities (Angelaki, Gu, & DeAngelis, 2009; De Gelder & Bertelson, 2003; Vines, Krumhansl, Wanderley, & Levitin, 2006). Yet, cross-sensory adaptation is not as well studied or understood. When different sensory modalities measure and respond to the same external stimulus, cross-sensory adaptation could enable cross-modal sharing of valuable statistical information regarding recent sensory events and context. The presence of cross-sensory adaptation might therefore expose functional adaption at the multimodal systems level – namely, higher-level brain processes that interpret multisensory information for perceptual decision-making.

Self-motion perception is an inherently multisensory function, relying on visual, vestibular, and other somatosensory cues (Greenlee, Frank, Kaliuzhna, Blanke, Bremmer, Churan, Cuturi, Macneilage, et al., 2016; Kaliuzhna, Ferrè, Herbelin, Blanke, & Haggard, 2016). Furthermore, self-motion perception demonstrates a high degree of adaptation to perturbations of environmental dynamics, e.g., at sea or in space (Duh, Parker, Philips, & Furness, 2004; Nachum et al., 2004; Shupak & Gordon, 2006). Accordingly, the adaptation of self-motion perception likely harnesses multisensory processes, applied on a continual and ongoing basis in normal brain function (Carriot, Jamali, & Cullen, 2015; Zaidel, Ma, & Angelaki, 2013; Zaidel, Turner, & Angelaki, 2011). However, our understanding of how cross-sensory adaptation affects self-motion perception is limited.

Crane et al. (2013) found that short-duration (1.5s) self-motion stimuli (either visual or vestibular) do not cause cross-sensory adaptation. Subsequently, Cuturi and MacNeilage (2014) demonstrated that longer duration visual self-motion stimuli (15s) do cause vestibular adaptation, concluding that cross-sensory (visual→vestibular) adaptation only occurs after exposure to visual stimuli presented for at least 15s. However, Cuturi and MacNeilage (2014) did not test the adaptive effects of vestibular stimuli on visual perception (vestibular→visual). Also, both studies tested the effect of a single ‘adapting’ stimulus. It is currently unknown how stimulus event history, beyond a single stimulus, affects cross-modal perception. Here, we hypothesized that experiencing several short-duration (1s) stimuli might elicit cross-sensory adaptation (even when their sum, if presented together, would be too short to elicit cross-sensory adaptation).

To address this question, we devised an experiment that tested the cross-modal effects of experiencing several biased (short duration, 1s) visual or vestibular self-motion stimuli. Trials were grouped in 140 batches, each comprising three ‘prior’ trials, with headings biased to the right or left, followed by a single unbiased ‘test’ trial. Right and left biased batches were interleaved pseudo-randomly. The experiment included four conditions: two uni-sensory (using only visual or only vestibular stimuli) and two cross-sensory (testing the effects of vestibular heading discrimination on subsequent visual perception, and vice versa). Not surprisingly, significant adaptation was seen in both unisensory conditions. Strikingly, we found significant adaptation also in both cross-sensory conditions, even with 1s duration stimuli. By fitting a logistic regression model to dissociate the effects of prior stimuli and prior choices, we found that prior stimuli (not choices) accounted for the adaptation of heading perception. These results indicate that the brain monitors and adapts to the statistics of stimulus events, cross-modally.

## METHODS

### Participants

Forty-five healthy adults participated in this study (21 male, 24 female; mean age ± SD = 23.8 ± 2.8 years, age range: 18 – 33). This study was approved by the Bar Ilan University, Gonda Brain Research Center Ethics Committee, and all participants signed informed consent. Participants were recruited via online student forums and received monetary compensation or course credit in lieu of participation. All participants had normal or corrected to normal vision and no reported neurological condition or chronic medication.

Participants were recruited to perform one of the four experimental conditions, which was randomly chosen. Then they were invited to perform the rest of the conditions (performed on four different days). We did not require that each participant perform all four conditions (this was not a within-subject analysis). Of the 45 participants, eight performed four conditions, five performed three conditions, one participant performed two conditions, and 31 participants performed one condition. Each condition (four independent datasets) comprised 20 participants.

### Stimuli and task

The participants were seated comfortably in a car seat mounted on a motion platform (MB-E-6DOF/12/1000, Moog Inc.), and restrained safely in the seat with a 4-point harness. Head movements were limited by a head support (Black bear, Matrix Seating Ltd.). Vestibular motion cues comprised inertial translations of the motion platform. Visual motion cues simulated self-motion through a 3D star field (optic flow) and were presented using a head-mounted display (HMD; Oculus Rift) worn by the participants. Figure 1A depicts a participant in the experimental setting (lights on only for the picture; the lights were off during the experiment).

**Figure 1.**
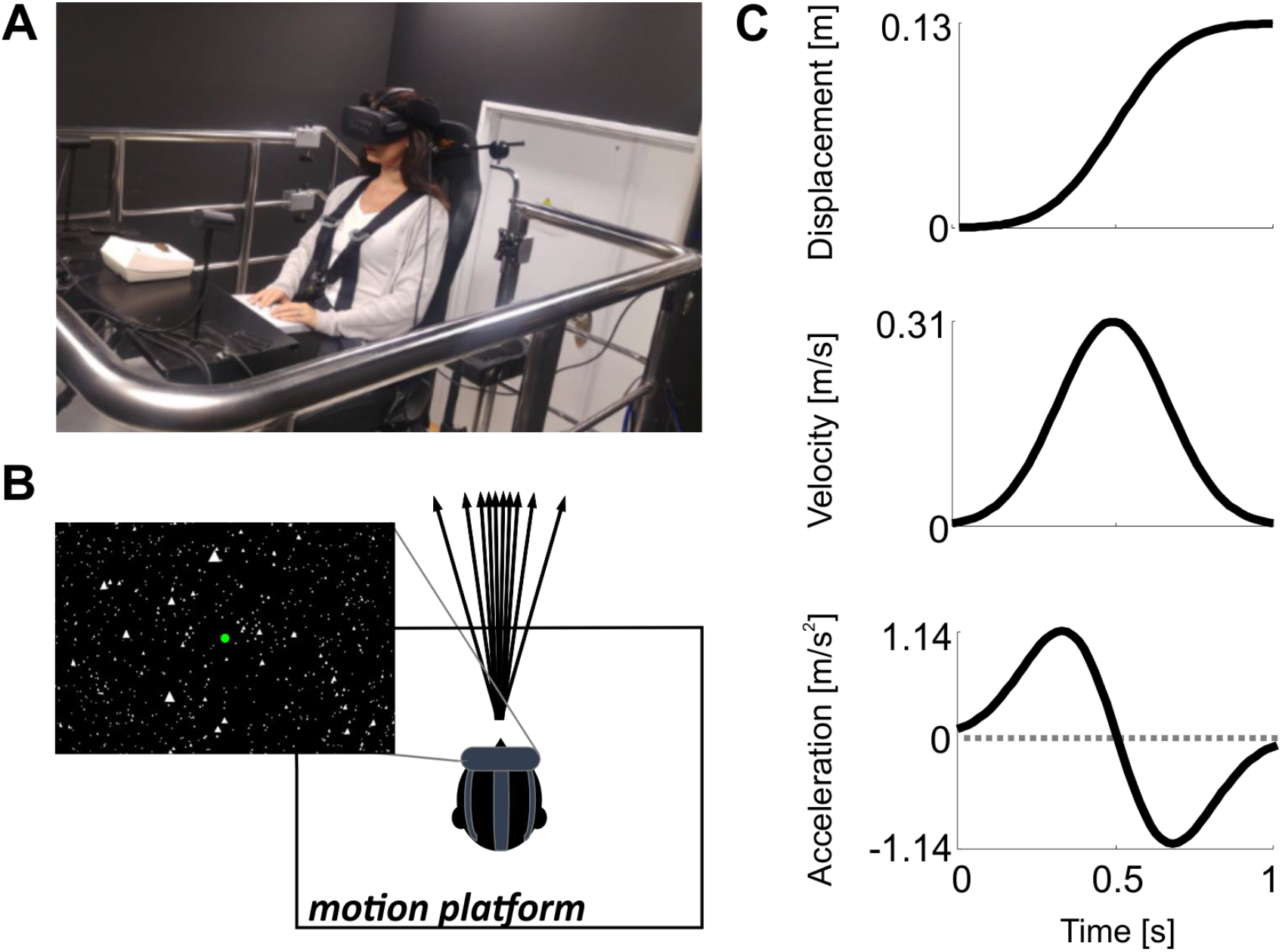
Experimental paradigm. (A) A participant seated in the multisensory motion simulator (lights on only for the picture; the lights were off during the experiment). (B) Schematic illustration of the motion platform from above. Vestibular stimuli comprised inertial motion of the motion platform. Visual stimuli (inset) simulated self-motion through a 3D star field (optic flow), and were presented in virtual reality via a head-mounted display. Arrows depict the visual and vestibular self-motion heading directions. (C) Motion profiles of the (visual and vestibular) self-motion stimuli.

The vestibular and visual (simulated) self-motion stimuli comprised single-interval linear motions, in a primarily forward-moving direction, with slight deviations to the right or left of straight ahead (Fig. 1B). The stimuli followed a Gaussian velocity motion profile with 1s duration and total displacement 13 cm (peak velocity 0.31 m/s and peak acceleration 1.14 m/s^2^; Fig. 1C).

The participant’s task was to discriminate whether their perceived heading (for each stimulus presentation) was to the right or left of straight-ahead (two-alternative forced-choice). Stimuli were self-initiated by pressing a central “start” button on the response box (Cedrus RB-540) and choices were reported, after the stimulus had ended, by pressing the corresponding right or left button on the response box. The participants were instructed to focus on a central fixation point, which remained present throughout the duration of the experiment. Different auditory signals were used to indicate: i) when the system was ready for trial initiation, ii) that a choice selection was registered, or iii) choice time-out (2s after the stimulus had ended; participants were instructed to avoid this by always making a choice and to make the best guess when unsure). No feedback about correct or incorrect choices was provided.

### Experimental conditions and trial structure

To investigate how the prior stimuli biased self-motion perception both within and across modalities, we tested four experimental conditions: 1) only vestibular stimuli were presented. In this condition, we tested the effects of prior vestibular heading discriminations on subsequent vestibular perception. Accordingly, both the ‘adaptor’ and the ‘test’ stimulus were vestibular (abbreviated: ves→ves); 2) only visual stimuli were presented. In this condition, we tested the effects of prior visual heading discriminations on subsequent visual perception. Accordingly, both the adaptor and the test stimulus were visual (abbreviated: vis→vis); 3) visual and vestibular stimuli were presented. In this condition, we tested the effects of prior visual heading discriminations on subsequent vestibular perception. Accordingly, the adaptor stimulus was visual, while the test stimulus was vestibular (abbreviated: vis→ves); and 4) vestibular and visual stimuli were presented. In this condition, we tested the effects of prior vestibular heading discriminations on subsequent visual perception. Accordingly, the adaptor stimulus was vestibular, while the test stimulus was visual (abbreviated: ves→vis).

In each condition, the trials were grouped into small ‘batches’ – each comprising a series of three biased ‘prior’ trials (with the adaptor stimulus), followed by a single unbiased ‘test’ trial with the test stimulus (Fig. 2A). Headings for the prior trials (*h_prior_*) were drawn from one of two normal distributions: *h_prior+_ ~N*(*μ* = +5°, *σ* = 2.5°) or *h_prior_*– *~N*(μ = –5°, *σ* = 2.5°) for rightward and leftward biased batches, respectively (where 0° represents straight ahead, and positive and negative values reflect rightward and leftward headings, respectively). In each experimental block, half of the batches were biased rightward and the other half leftward, interleaved pseudo-randomly.

**Figure 2.**
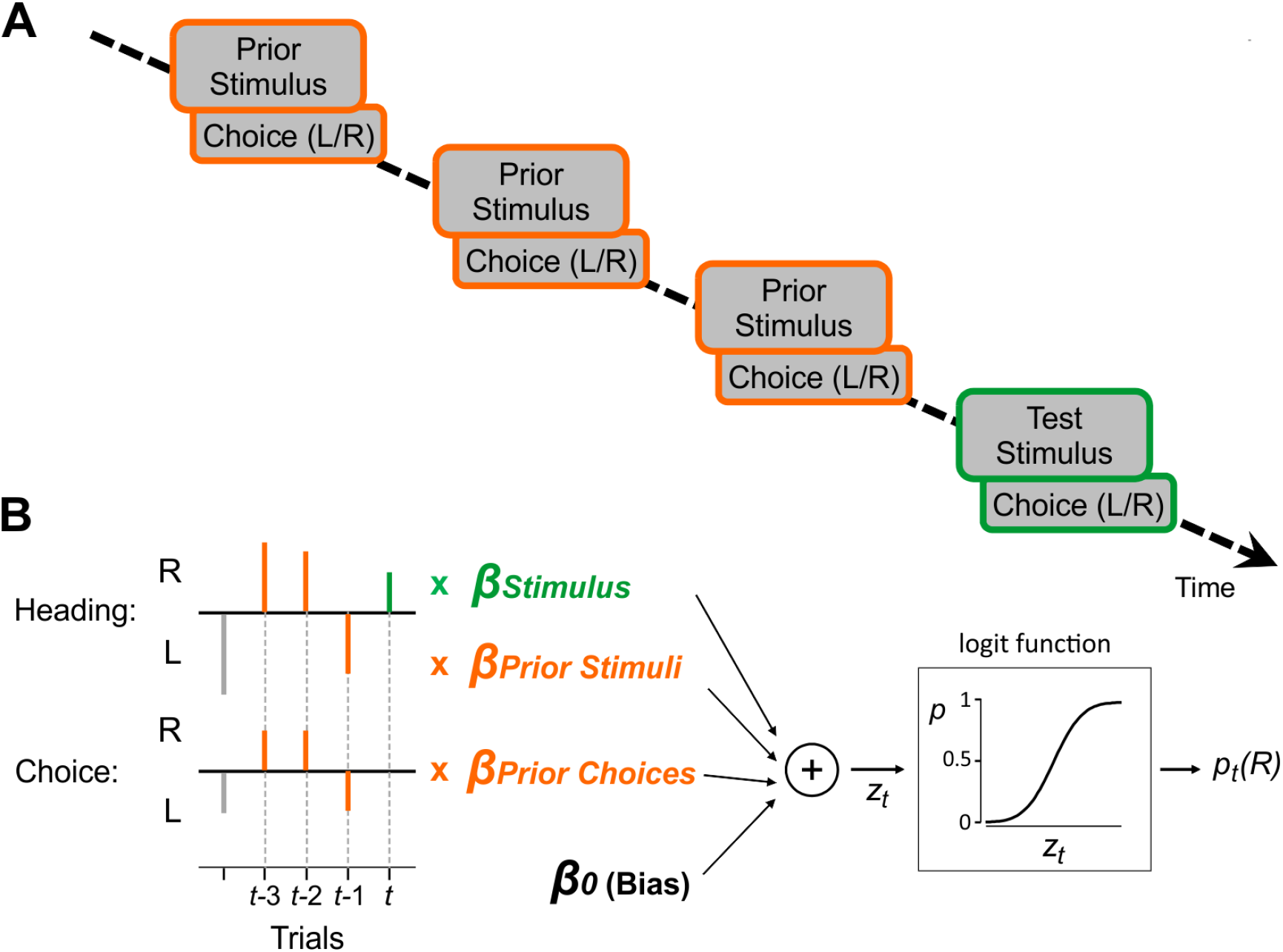
Experimental procedure and model. (A) A ‘batch’ of trials comprised three ‘prior’ trials (orange frames) followed by one ‘test’ trial (green frame). For each trial, the participants were required to discriminate whether the stimulus heading was to the right or left of straight ahead. The prior trials in a batch were biased either to the right or to the left. Rightward and leftward biased batches were interleaved in a pseudorandom manner (70 batches each, 140 in total). (B) A binomial logistic regression (fit per participant) predicted the choice on trial t by a weighted linear sum (z_t_) of four predictors: the current trial’s heading direction (green stem, × β_Stimulus_), average heading direction of the three prior trials (orange ‘heading’ stems, × β_Prior.Stimuli_), mode of the three prior choices (orange ‘choice’ stems, × β_Prior.Choices_) and general bias (β_0_). The sum (z_t_) was passed through a logit function to calculate the probability of making a rightward choice, p_t_(R) on trial t.

For test trials, the heading sign was selected randomly (50% probability for right or left) and heading magnitude followed a staircase procedure (Cornsweet, 1962). Possible heading values were spaced logarithmically around 0°: *h_test_* = ±16°, ±8°, ±4°, ±2°, ±1°, ±0.5°, ±0.25°. The staircase started at the easiest heading magnitude (|*h_test_*| = 16°). After a correct response, heading magnitude was reduced (i.e., became more difficult) with 30% probability and remained unchanged for the remaining 70%. After an incorrect response, it was increased (i.e., became easier) with 80% probability and remained unchanged for the remaining 20%. This staircase rule converges to ~73% correct responses (MacNeilage, Banks, DeAngelis, & Angelaki, 2010) – a highly informative region of the psychometric function. A separate staircase was run for the test trials from the leftward and rightward biased batches. Participants were instructed to discriminate the heading direction for all stimuli presented and were not informed about any trial structure (i.e., our definition of prior and test trials). Therefore, to the participants, the stimuli appeared as a continuous stream of trials. When questioned after the experiment, none reported noticing any trial structure.

An experimental session comprised 140 trial batches (70 biased to each side). To improve alertness and attention, the session was divided into two equal blocks with a short break in between. These were later merged for analysis. To speed up the convergence of the staircase, at the beginning of each block 10 additional test trials without priors were presented for each staircase. These were included in the psychometric fits, but not in the logistic regression model fits (which required ‘prior’ trials for the model inputs).

### Data analysis

Data analyses and statistics were performed using Matlab R2013b (MathWorks), and the psignifit toolbox for Matlab (version 4; Schütt, Harmeling, Macke, & Wichmann, 2016). For further details on statistical comparisons, see the *Statistics* section below. Psychometric plots were defined as the proportion of rightward choices as a function of heading angle (*h*) and calculated by fitting the data with the psychometric function:

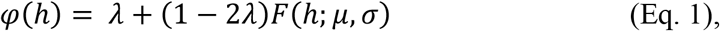

where *F*(*h*; *μ*, *σ*) is a cumulative Gaussian distribution function with mean = *μ* and standard deviation (SD) = *σ* (Klein, 2001). In terms of behavior, *μ* represents the point of subjective equality (PSE) which is the heading estimate for which the participant would choose right/left with equal probability (also known as the ‘bias’), *σ* represents the psychophysical threshold (lower values reflect better performance), and *λ* represents the lapse rate, which is the rate of reporting an incorrect choice even for obvious stimuli (assumed symmetrical for right and left). Two psychometric functions were computed (per condition) using data only from the test trials, separated according to the priors’ bias (left/right). To quantify the behavioral effect of the priors on perception, the difference between the PSEs of these two psychometric functions was calculated (per condition):

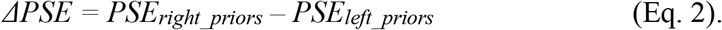

Here, positive ΔPSE values mean that prior trials had a repulsive effect on subsequent perception. Namely, after experiencing rightward biased stimuli, the probability of choosing left for the test stimulus was increased (and vice versa for leftward biased stimuli). We refer to this (repulsive) shift as ‘adaptation’ because it is in the same direction as other effects termed adaptation (such as motion aftereffects; Anstis et al., 1998; Gibson, 1937; Gibson & Radner, 1937; Thompson, P., & Burr, 2009). By contrast, negative ΔPSE values reflect an attractive effect of prior stimuli. Namely, following rightward biased stimuli, the probability of choosing right for the test stimulus was increased (and vice versa for leftward biased stimuli). This attractive shift is often termed a ‘repetition’ or ‘consistency’ bias (Alexi et al., 2018; Fischer & Whitney, 2014; Liberman, Fischer, & Whitney, 2014; Liberman, Manassi, & Whitney, 2018; Taubert & Alais, 2016).

### Probabilistic choice model

The measure of ΔPSE (Eq. 2) quantifies the overall effects of biased prior trials on behavior. However, it does not dissociate whether these effects arise due to the prior stimuli, prior choices, or a combination thereof. To dissociate the separate effects of prior stimuli and prior choices on subsequent performance, we adapted the probabilistic choice model from Busse et al. (2011), Abrahamyan et al. (2016), and Feigin et al (2021) to our experiment (Fig. 2B). According to the model, the probability to choose rightward on a given test trial *t* follows the binomial logistic regression:

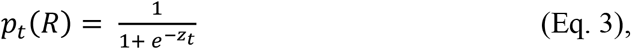

where *z_t_* is the log-odds ratio of the probability to choose rightward or leftward on a specific test trial, calculated by a linear combination of predictors:

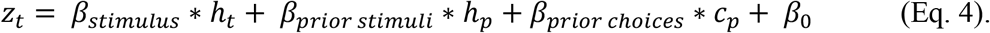

Here, *h_t_* is the heading direction on the current test trial, *h_p_* is the average heading direction of the three preceding prior trials, and *c_p_* is the mode of the choices from the three preceding prior trials (we also tested the model using the average choice, described further below). Choices were represented by –1 or +1 for leftward and rightward choices, respectively. Heading directions were normalized by the root mean square (RMS) such that both heading and choice predictors had RMS = 1 (to make these parameters comparable). The regression weights *β_stimulus_, β_prior_stimuli_, β_prior_choices_*, and baseline bias *β_0_* were fit per participant and condition.

To assess the effects of the predictors on the perceptual choices, we considered four models: 1) M_0_: “no history” model – this is the most basic model that does not take prior stimuli or prior choices into account (*β_prior_stimuli_* and *β_prior_choices_* from Eq. 4 are set to zero). It therefore includes only the baseline bias and current stimulus intensity as predictors. 2) M_1_: “prior stimuli” model – takes the prior stimuli (but not prior choices) into account (*β_prior_choices_* from Eq. 4 is set to zero). 3) M_2_: “prior choices” model – takes the prior choices (but not the prior stimuli) into account (*β_prior_stimuli_* from Eq. 4 is set to zero). 4) M_3_: “prior choices + prior stimuli” model – takes into account all the predictors from Equation 4, including the prior stimuli and prior choices (no coefficients set to zero).

The results (presented below) revealed that prior stimuli (and not prior choices) had a significant effect on the participant’s choices. Thus, to further investigate the effects of stimulus history we also fit an expanded model based on M_3_ (M_3E_: “prior choices + separate prior stimuli” model) that took into account each stimulus (individually) from the three prior trials, such that the linear combination of predictors was:

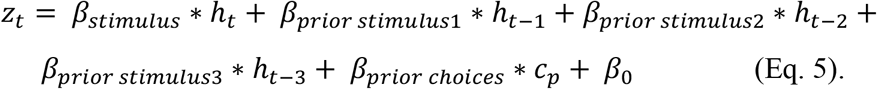

Here, *h_t-1,_ h_t-2_*, and *h_t-3_* are the heading directions of the three prior trials, where *t-*1 represents the trial immediately preceding the test trial*, t-*2 the trial before *t-*1, and *t-*3 the trial before *t-*2 (the first prior in the batch).

### Parameter recovery

To verify that the model fits for M_3E_ were plausible and unbiased, we simulated data using the beta parameters from the model fits and checked that these were reliably recovered. We present simulations of the vis-vis condition (Suppl. Figs. 2–3) because its results showed a graded effect of *β_prior_stimulus_* (from *t*-1 to *t*-3), providing simple expectations against which to compare the simulated results. Parameter recovery using the other conditions worked just as well. Simulations were performed for each one of the 20 participants using their beta parameters and the logistic regression model (Eqs. 3 and 5) to simulate ‘choices’.

Heading directions for the simulated prior trials were drawn from the same prior distributions as those used in the experiments, and simulated ‘test’ trial headings followed the same staircase procedure as the experiments (described in the *Experimental conditions and trial structure* section above). For the first three (prior) trials of the simulation, responses were simulated based only on the stimulus (*β_stimulus_*) and general bias (*β_0_*). From trial four (the first test trial) onwards, responses for all trials (test and prior) were simulated in the following way: the linear combination of the predictors (Eq. 5) was computed using the participant’s six beta parameters from M_3E_ (‘original betas’). This was used to compute the odds for a rightward choice on the current trial (Eq. 3). A binomial value (‘1’ for right and ‘-1’ for left) was then randomly drawn according to the computed odds. Each response was marked as ‘correct’ or ‘incorrect’ according to the sign of the heading stimulus (for the staircase). The simulated choices for the test trials were then fit (in the same way that the participants’ data were fit) to extract the six betas again (‘recovered betas’). This process was repeated 100 times, resulting in 100 sets of 6 recovered betas per participant. We then compared the median of each recovered beta to the participant’s original betas.

In addition, we also simulated a fictitious participant whose “original” betas were set by us. This was done to better visualize and to further test the model fit. We used three representative and different (equispaced) values for the three prior stimuli betas (*β_prior_stimulus1_* = −0.2, *β_prior_stimulus2_* = −0.1, *β_prior_stimulus3_* = 0) to confirm that these were accurately and individually recovered (also when simulated in reverse order, described further below). The other beta parameters were set to: *β_stimulus_* = 0.9, *β_prior_choices_* = 0.3 and *β_0_* = 1.5 (to test that a bias was accurately recovered). Simulating with other combinations of betas also resulted in full recovery of the beta values. To further test that the model fitting was robust to different compositions of prior stimulus coefficients, we also simulated and recovered the beta coefficients after reversing the order of the original *β_prior_stimulus1,_ β_prior_stimulus2_*, and *β_prior_stimulus3_* as input (i.e., the original *β_prior_stimulus3_* was switched with *β_prior_stimulus1_*; Suppl. Fig. 3).

### Model comparisons

We used Bayesian model comparisons to compare M_0_, M_1_, M_2,_ and M_3_ (which had different numbers of parameters), and thereby assess the added value of taking into account stimulus and choice information from prior trials. This was done by computing the Akaike information criterion (AIC) score for each model:

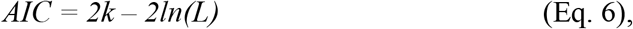

where *k* is the number of parameters in the model and *L* is the maximum value of the likelihood function for the model. We then calculated the difference between the models’ AIC scores:

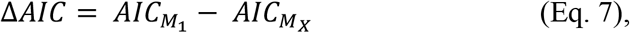

where M_*X*_ represents a particular model (i.e., M_0_, M_2,_ or M_3_) being compared to M_1_, which was used as the basis for comparison. We chose the M_1_ (“prior stimuli” model) as the basis for the comparison based on the results for M_3_ (below) that prior stimuli (and not prior choices) were significant predictors. We then used ΔAIC to calculate the Bayes factors (BFs) as follows:

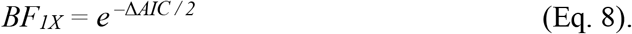

BF_1X_ > 1 indicates a preference for M_1_ over the alternative model M_*X*_, and BF_1X_ < 1 indicates a preference for the alternative model.

In addition to AIC we also computed the Bayesian information criterion (BIC), which penalizes the model complexity more heavily:

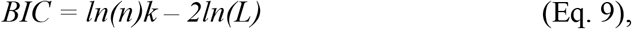

where *k* and *L* are the same parameters from Equation 6, and *n* is the number of observations.

### Statistics

Our apriori hypothesis for this experiment was that biased ‘prior’ trials would lead to an adaptation aftereffect on subsequent ‘test’ trials. Four sets of experimental data were gathered to test this (two uni-sensory conditions and two cross-sensory conditions). For each dataset, we tested whether ΔPSE (Eq. 2) was significantly different from zero, using a two-tailed *t*-test. Since these were four independent datasets (each with the same hypothesis), we did not adjust the resulting p-values.

Our subsequent investigation into the source of the adaptation aftereffect compared the model parameters *β_prior_stimuli_* and *β_prior_choices_* (Eq. 4) to zero using two-tailed *t*-tests. When investigating the significance of these parameters we multiplied the raw *p*-values by 2 (Bonferroni correction for multiple comparisons) and present 97.5% (instead of 95%) confidence intervals. Although we also compared the effects of the current stimulus (*β_stimulus_*) and general bias (*β_0_*) – these comparisons were only a sanity check to confirm (trivially) that the current stimulus had a strong influence on perceptual choices, and that there was no unexpected baseline bias.

The last analysis (M_3E_) was more exploratory in nature and aimed to further investigate our finding that prior stimuli affected subsequent perception. Specifically, we analyzed the individual contributions of each of the three prior stimuli. Due to the exploratory nature of this analysis, we present the raw *t*-test results (uncorrected), interpret these results with caution, and call for further research to confirm and to better understand them.

## RESULTS

In this study, we investigated cross-modal adaptation of visual (“vis”) and vestibular (“ves”) self-motion perception. Specifically, we tested whether several short-duration (1s) ‘prior’ stimuli (with biased headings) from one cue would bias subsequent heading perception of the other cue. Four conditions of different adaptor and test stimuli were run (abbreviated: adaptor→test): two uni-sensory (vis→vis and ves→ves), and two cross-sensory (vis→ves and ves→vis).

### Cross-sensory adaptation of self-motion perception

To expose adaptation, two psychometric curves (one for each prior bias – to the right and left) were generated per participant and condition. Responses only to the test stimuli were used to create these plots. If the prior trials did not affect subsequent choices, then the two psychometric curves should lie roughly on top of each other and have the same point of subjective equality (PSE). However, if the prior trials biased subsequent choices, then the psychometric curves should appear shifted relative to one another.

Example psychometric functions for the four conditions are presented in Figure 3A. A difference in PSE between the two psychometric functions can be seen in all four conditions: both in the uni-sensory and cross-sensory conditions (Fig. 3A, left and right columns, respectively). More specifically, these shifts reflect a repulsive bias (adaptation). Namely, an increased probability for rightward choices following leftward biased priors (and vice versa). This is seen by a leftward shift (meaning more rightward choices) for the curves with left priors (lighter shades) and a rightward shift (meaning more leftward choices) for the curves with right priors (darker shades). To quantify this, we calculated the difference between the two psychometric curves’ PSEs (ΔPSE, Eq. 2) per participant, and condition.

**Figure 3.**
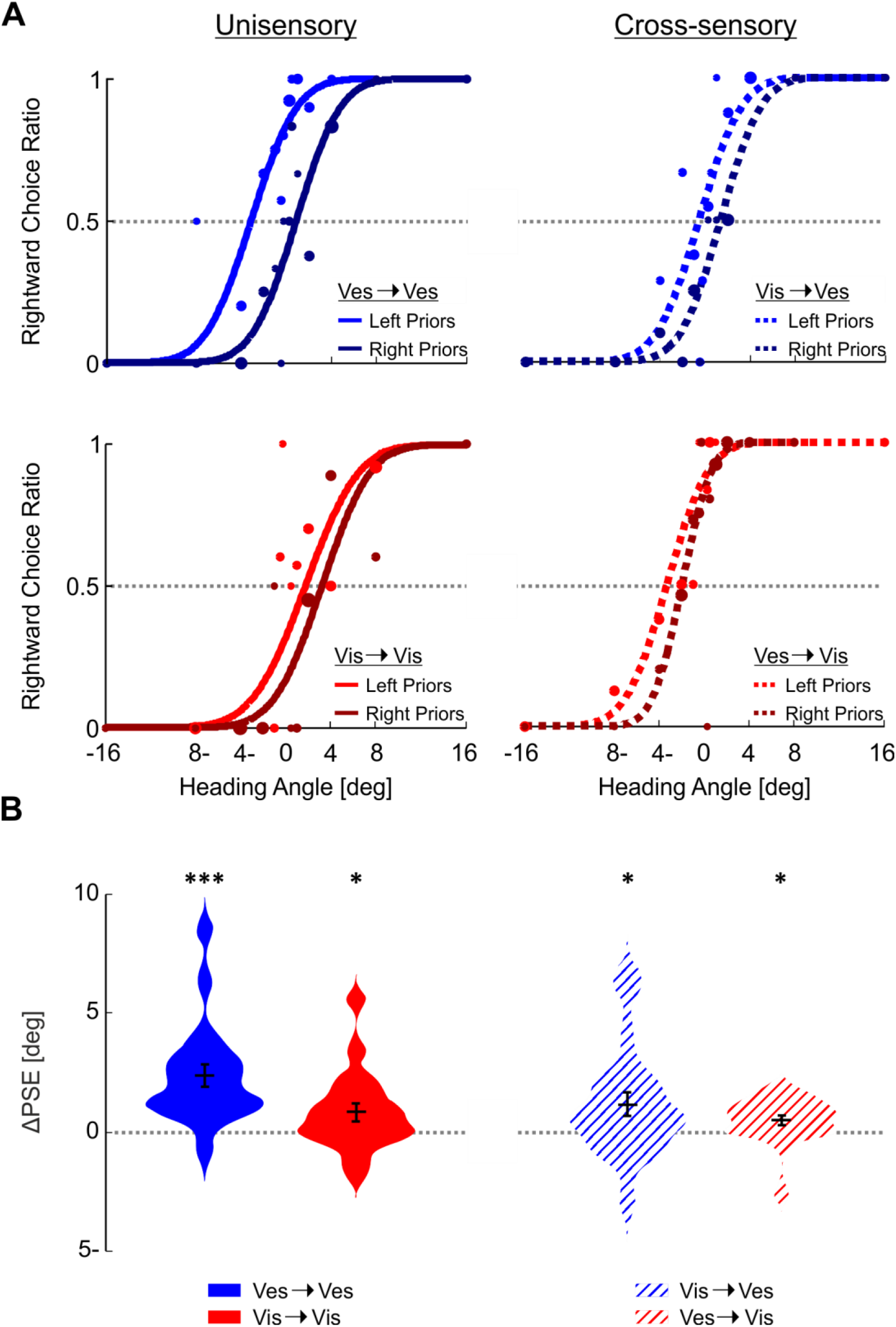
Psychometric shifts. (A) Example psychometric curves calculated from responses to the test trials (only), for the four different conditions (subplots). Blue and red colors represent responses to vestibular and visual test trials, respectively. For each condition, two separate psychometric curves were calculated according to the prior type (left or right biased priors, represented by light and dark colors, respectively). The data (circles) present the ratio of rightward choices per stimulus heading direction (and prior type), and were fit with cumulative Gaussian distribution functions. The circle sizes reflect the number of trials for a given stimulus heading and prior type. Solid and dashed lines represent uni-sensory and cross-sensory conditions, respectively. (B) Summarized (group) PSE difference (ΔPSE = PSE after rightward biased priors minus PSE after leftward biased priors). \*\*\**p* < 0.001, \**p* < 0.05.

At the group level, significant adaptation (positive ΔPSE shifts) was seen for both of the unisensory conditions, as well as both of the cross-sensory conditions (Fig. 3B; ves→ves: *t*_19_ = 4.92, ***p* = 9.6∙10^−5^**, *Cohen’s d* = 1.10, 95% *CI* = [1.33 3.30]; vis→vis: *t*_19_ = 2.27, ***p* = 0.035**, *Cohen’s d* = 0.51, 95% *CI* = [0.06 1.60]; vis→ves: *t*_19_ = 2.26, ***p* = 0.035**, *Cohen’s d* = 0.51, 95% *CI* = [0.09 2.24]; ves→vis: *t*_19_ = 2.35, ***p* = 0.029**, *Cohen’s d* = 0.53, 95% *CI* = [0.05 0.93]; *t*-tests). These results indicate that cross-sensory adaptation occurs after experiencing several short-duration stimuli.

### Stimulus (not choice) adaptation

We have shown above that trials with biased stimuli lead to subsequent adaptation (both within and across modalities). But, does this result from the biased stimuli per se or perhaps from the choices? Although choices and stimuli are correlated, they are not perfectly correlated (and are therefore separable) for two reasons: i) choices are binary whereas stimuli are continuous, and ii) participants naturally make mistakes.

To investigate whether the observed adaptation was stimulus-related or choice-related, we fitted all the trials’ choices with a logistic regression model (using the “prior choices + prior stimuli” model, M_3_; see Fig. 2B and Methods for model details) with four predictors: *h_t_* (current stimulus heading), *h_p_* (prior stimuli headings), *c_p_* (prior choices) and general subjective bias (Eqs. 3 and 4). This enabled us to separate the effects of prior stimuli from prior choices, by their respective, fitted weights.

Model fits revealed that the prior stimuli (and not prior choices) accounted for the adaptation. The prior stimuli weights (*β_prior_stimuli_*) were significantly negative for both uni-sensory conditions as well as both cross-sensory conditions (Fig. 4, second row: ves→ves: *t*_17_ = −5.76, ***p* = 4.6∙10^−5^**, *Cohen’s d* = −1.36, 97.5% *CI* = [−1.97 −0.77]; vis→vis: *t*_19_ = −5.14, ***p* = 1.2∙10^−4^**, *Cohen’s d* = −1.15, 97.5% *CI* = [−1.50 −0.52]; vis→ves: *t*_19_ = −2.93, ***p* = 0.018**, *Cohen’s d* = −0.66, 97.5% *CI* = [−0.94 −0.08]; ves→vis: *t*_19_ = - 2.85, ***p* = 0.020**, *Cohen’s d* = −0.64, 97.5% *CI* = [−0.65 −0.04]; *t*-tests; *p*-values reported here were multiplied by two to correct for multiple comparisons). Negative weights here indicate a repulsive effect (adaptation). By contrast, the prior choices weights (*β_prior_choices_*) were not significantly different from zero in three out of the four conditions (Fig. 4, third row: vis→vis: *t*_19_ = 1.54, ***p* = 0.28**, *Cohen’s d* = 0.34, 97.5% *CI* = [−0.17 0.71]; vis→ves: *t*_19_ = 0.68, ***p* > 1**, *Cohen’s d* = 0.15, 97.5% *CI* = [−0.28 0.49]; ves→vis: *t*_19_ = 1.02, ***p* = 0.64**, *Cohen’s d* = 0.23, 97.5% *CI* = [−0.18 0.43]; *t*-tests; *p*-values reported here were multiplied by two to correct for multiple comparisons), and significantly positive (i.e., ‘attractive’, in the opposite direction to adaptation) for the ves→ves condition (*t*_17_ = 3.28, ***p* = 8.8∙10^−3^**, *Cohen’s d* = 0.77, 97.5% *CI* = [0.12 0.87]; *t*-test; the *p*-value reported here was multiplied by two to correct for multiple comparisons).

**Figure 4.**
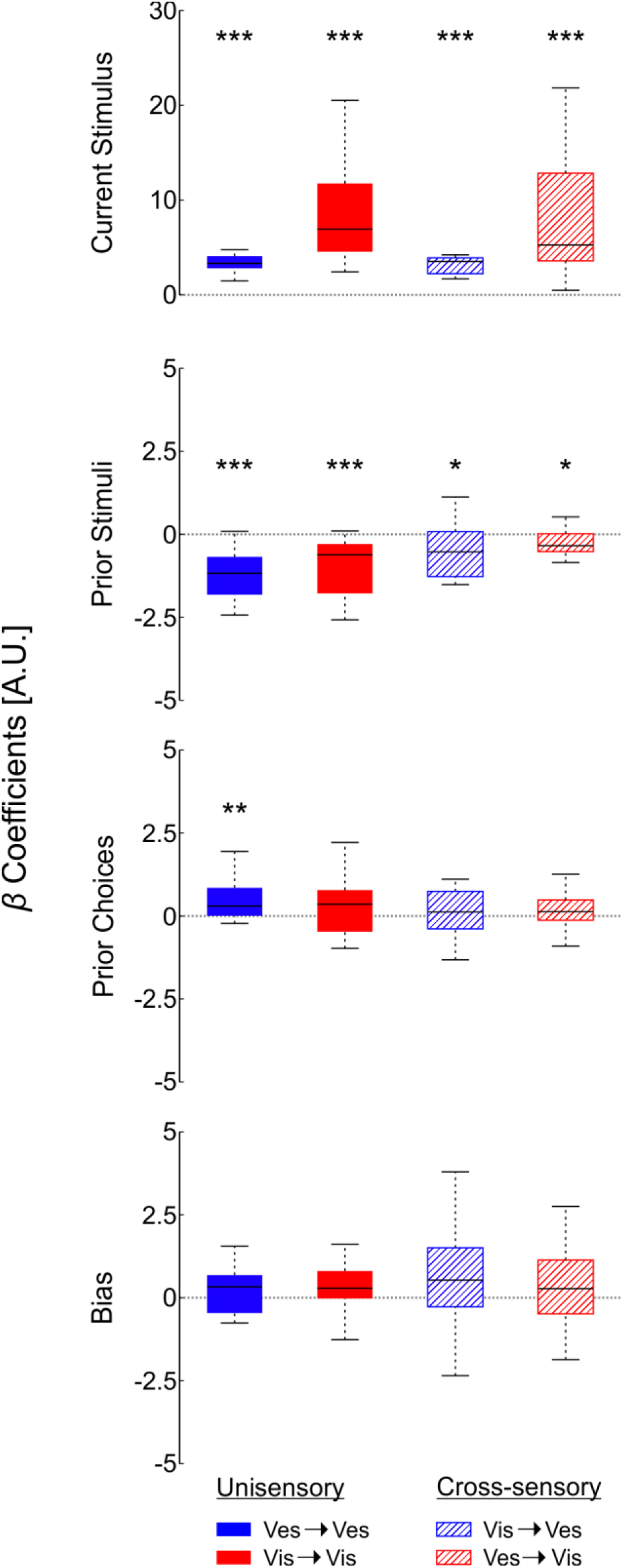
M_3_ (“Pror choices + prior stimuli” model) results summary. Beta coefficient values for the four model parameters (rows) per condition. Blue and red colors reflect conditions with vestibular and visual test trials (respectively), and filled and striped textures reflect uni-sensory and cross-sensory conditions (respectively). Black horizontal lines inside the boxes represent the medians. Upper and lower box limits mark the interquartile ranges. \*\*\**p* < 0.001, \*\**p* < 0.01, \**p* < 0.05.

These results indicate that adaptation of heading perception (visual, vestibular, and cross-sensory) is specifically to the prior stimuli (and not to the choices themselves). When an effect of prior choices was present (ves→ves), it was in the opposite direction to the stimulus effect (and to the overall PSE shifts, which represent an aggregate/combined effect of prior stimuli and prior choices).

Naturally, the current stimulus predictor had the largest weight (*β_stimulus_*) and was significantly positive in all conditions (Fig. 4, top row: ves→ves: *t*_17_ = 10.09, ***p* = 1.4∙10^−8^**, *Cohen’s d* = 2.38, 95% *CI* = [2.89 4.41]; vis→vis: *t*_19_ = 8.11, ***p* = 1.4∙10^−7^**, *Cohen’s d* = 1.81, 95% *CI* = [6.39 10.84]; vis→ves: *t*_19_ = 9.25, ***p* = 1.8∙10^−8^**, *Cohen’s d* = 2.07, 95% *CI* = [2.69 4.26]; ves→vis: *t*_19_ = 5.40, ***p* = 3.3∙10^−5^**, *Cohen’s d* = 1.21, 95% *CI* = [4.86 11.01]; *t*-tests). This confirms that the current stimulus heading was the most crucial factor in forming the perceptual decisions. The general bias weight (*β_0_*) was not significantly different from zero in any condition (Fig. 4, bottom row: ves→ves: *t*_17_ = 1.21, ***p* = 0.24**, *Cohen’s d* = 0.29, 95% *CI* = [−0.14 0.51]; vis→vis: t_19_ = 1.47, ***p* = 0.16**, *Cohen’s d* = 0.33, 95% *CI* = [−0.17 0.98]; vis→ves: *t*_19_ = 2.00, ***p* = 0.060***Cohen’s d* = 0.45, 95% *CI* = [−0.03 1.40]; ves→vis: *t*_19_ = 1.66, ***p* = 0.11**, *Cohen’s d* = 0.37, 95% *CI* = [−0.11 0.98]; *t*-tests). We also fitted the model with the prior choices parameter (*c_p_* from Eq. 4) taken as the average of the three prior choices (instead of the mode). Results were qualitatively similar (with only minor quantitative differences).

### Bayesian model comparisons

In the model analysis above, we found that the prior stimuli (and not the prior choices) accounted for the adaptation effect. Thus, for model comparison, we used the “prior stimuli” model (M_1_, which includes only the prior stimuli, current stimulus, and subjective bias parameters) as a base against which to compare the three other models. Bayes factors were calculated using the Akaike information criterion (AIC). Here, Bayes factors (BFs) greater than 1 (Fig. 5, bars to the right) indicate an advantage of M_1_ over the other model being compared. BFs > 3 (marked by a vertical dashed line in Fig. 5) provide substantial evidence in favor of M_1_ and BFs > 10 provide strong evidence (Jarosz & Wiley, 2014; Raftery, 1995; Wagenmakers, 2007).

**Figure 5.**
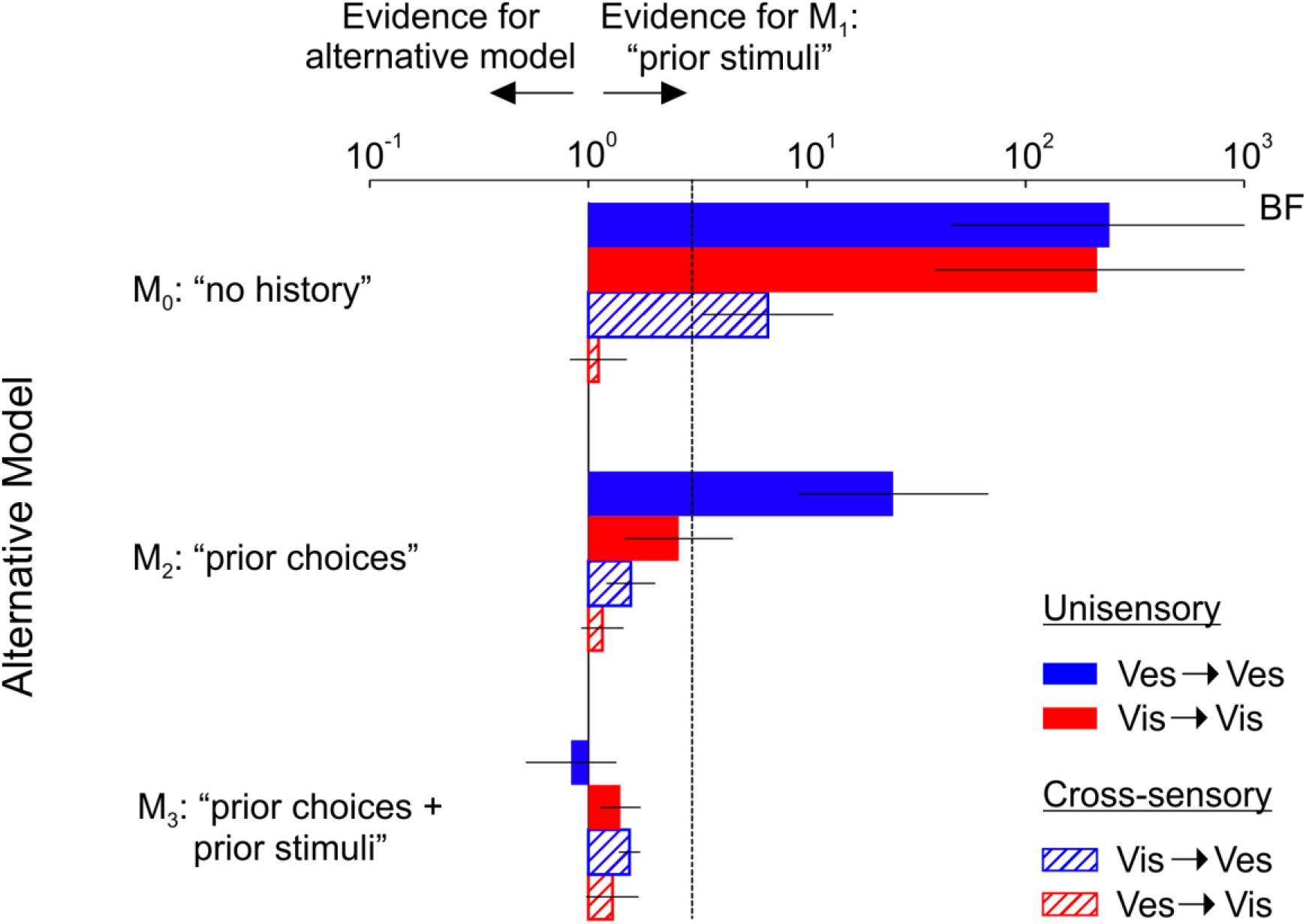
AIC Bayesian model comparisons. Bayes factors (mean ± SEM), calculated using the Akaike information criterion (AIC), reflect the likelihood of M1 vs. each of the alternative models (M0, M2 and M3). Values greater than one (bars to the right) indicate an advantage of M1 over the alternative model. Values greater than three (marked by a dashed line) indicate a substantial advantage of M1 over the alternative model. The same comparison using the Bayesian information criterion (BIC) is presented in Supplementary Figure 1.

The results indicate an advantage of M_1_ (“prior stimuli”) over M_0_ (“no history”) models (upper group of bars in Fig. 5) in the ves→ves (solid blue, mean BF = 242.48), vis→vis (solid red, mean BF = 212.92) and vis→ves (striped blue, mean BF = 6.62) conditions, but not in the ves→vis (striped red) condition, for which the models were comparable (mean BF = 1.11). This suggests that taking stimulus history into account improves the prediction of perceptual decisions enough to be a worthy component of the model in three out of the four conditions. A similar advantage was seen for M_1_ (“prior stimuli”) over M_2_ (“prior choices”; a middle group of bars in Fig. 5) for the ves→ves condition (mean BF = 24.80). Thus, even though a significant effect of prior choices was seen for ves→ves (Fig. 4), it was still better to use prior stimuli over prior choices when taking trial history into account. The other three conditions showed a weak preference for using prior stimuli over prior choices (mean BFs 1~3; possibly weakened by choice-stimulus correlations). M_1_ (“prior stimuli”) demonstrated a weak advantage over M_3_ (“prior choices + prior stimuli”; the bottom group of bars in Fig. 5) in three out of the four conditions (mean BFs 1~3). Together, these results suggest that it is justified to add prior stimuli, but not prior choices, to the model.

Comparisons using BIC, which penalizes the number of model parameters more strictly than AIC, show a clear advantage of taking history into account only for the unisensory (but not multisensory) conditions (Suppl. Fig. 1; M_1_ vs. M_0_). However, if taking prior trials into account, there is an advantage to using prior stimuli over prior choices (M_1_ vs. M_2_ and M_3_; bars to the right)

### Cross-sensory and uni-sensory adaptation are affected differently by prior stimuli

With the finding that adaptation in our paradigm was accounted for primarily by the prior stimuli (rather than prior choices), we further analyzed the individual contribution of each of the three prior stimuli. For this purpose, we fit the data with an expanded version of model M_3_ (M_3E_) that separated the individual effects of the three prior stimuli (Eq. 5) We note that this analysis was exploratory in nature (performed post-hoc given the results of this study) to identify trends in the data for further research in the future. Therefore, we interpret these results with caution.

Both uni-sensory conditions showed the same trend, and both cross-sensory conditions showed the same trend. Surprisingly, however, these trends were quite different. In the uni-sensory conditions adaptation seemed to be driven largely by the more recent heading stimuli (Fig. 6, top row, solid blue and red). Namely, those from trials *t-*1 (*β_prior_stimulus1_*: ves→ves: *t*_17_ = −4.27, ***p* = 5.1∙10^−4^**, *Cohen’s d* = −1.01, 95% *CI* = [−1.26 −0.43]; vis→vis: *t*_19_ = −3.44, ***p* = 2.8∙10^−3^**, *Cohen’s d* = −0.77, 95% *CI* = [−0.92 − 0.22]; *t*-tests) and *t*-2 (*β_prior_stimulus2_*: ves→ves: *t*_17_ = −6.35, ***p* = 7.3∙10^−6^**, *Cohen’s d* = - 1.50, 95% *CI* = [−0.73 −0.36]; vis→vis: *t*_19_ = −2.28, ***p* = 0.034**, *Cohen’s d* = −0.51, 95% *CI* = [−0.58 −0.03]; *t*-tests), and not *t-*3 (*β_prior_stimulus3_*: ves→ves: *t*_17_ = −0.32, ***p* = 0.75**, *Cohen’s d* = −0.08, 95% *CI* = [−0.33 0.24]; vis→vis: *t*_19_ = −1.54, ***p* = 0.14**, *Cohen’s d* = - 0.34, 95% *CI* = [−0.41 0.06]; *t*-tests).

**Figure 6.**
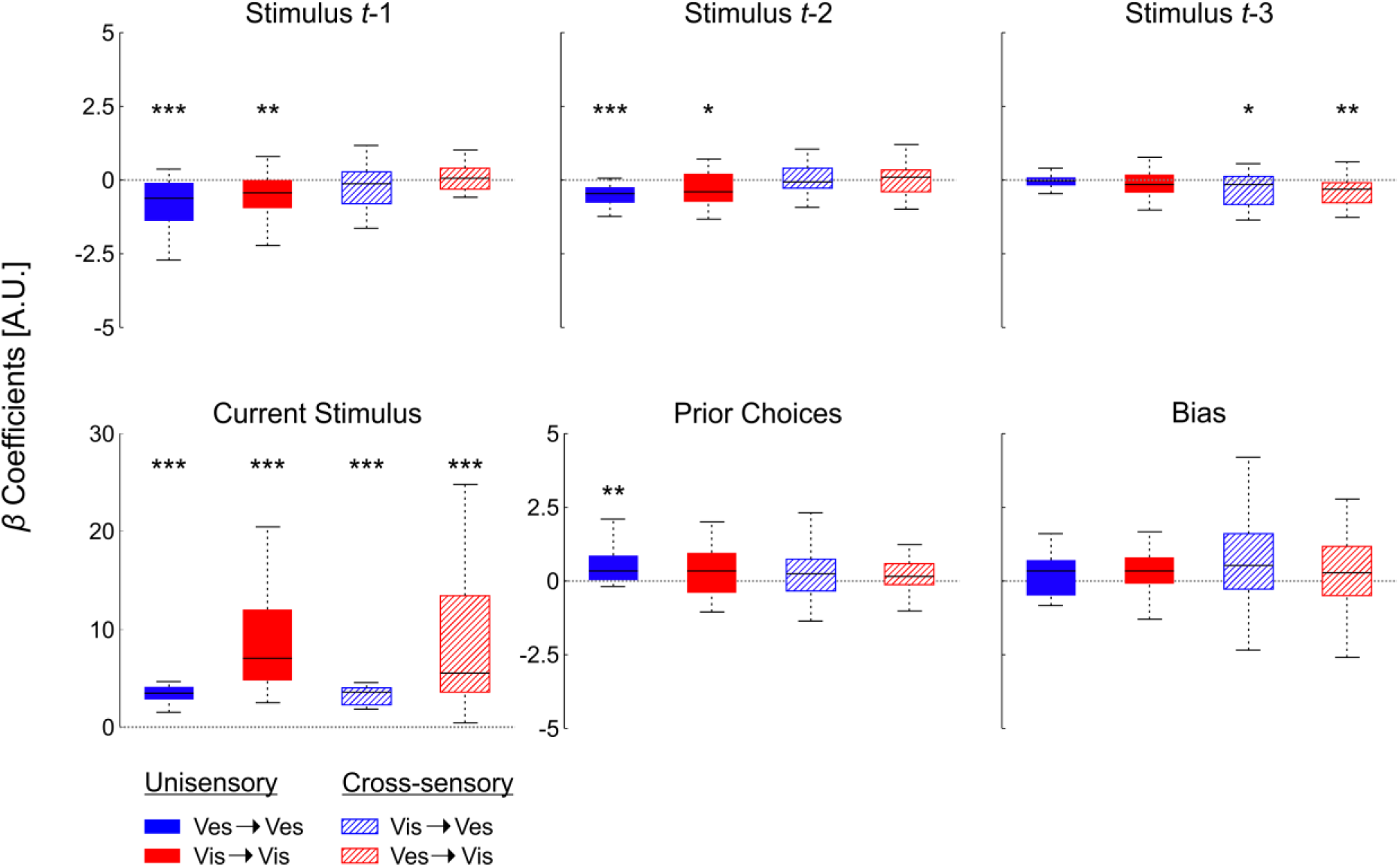
M_3E_ (“Prior choices + separate prior stimuli” model) results summary. Beta coefficient values for the six model parameters (subplots). Blue and red colors reflect conditions with vestibular and visual test trials (respectively), and filled and striped textures reflect uni-sensory and cross-sensory conditions (respectively). Black horizontal lines inside the boxes represent the medians. Upper and lower box limits mark the interquartile ranges. \*\*\**p* < 0.001, \*\**p* < 0.01, \**p* < 0.05.

By contrast, in the cross-sensory conditions, adaptation seemed to be influenced by the heading stimulus from trial *t-*3 (Fig. 6, top row, right plot: vis→ ves: *t*_19_ = −2.35, ***p* = 0.030**, *Cohen’s d* = −0.52, 95% *CI* = [−0.83 −0.05]; ves→vis: *t*_19_ = −2.95, ***p* = 8.2∙10^−3^**, *Cohen’s d* = −0.66, 95% *CI* = [−0.58 −0.10]; *t*-tests), but not from trials *t-*1 (left plot: vis→ves: *t*_19_ = −1.19, ***p* = 0.25**, *Cohen’s d* = −0.27, 95% *CI* = [−0.55 0.15]; ves→vis: *t*_19_ = 0.58, ***p* = 0.57**, *Cohen’s d* = 0.13, 95% *CI* = [−0.16 0.29]; *t*-tests) or *t-*2 (middle plot: vis→ves: *t*_19_ = 0.33, ***p* = 0.75**, *Cohen’s d* = 0.07, 95% *CI* = [−0.21 0.28]; ves→vis: *t*_19_ = −0.49, ***p* = 0.63**, *Cohen’s d* = −0.11, 95% *CI* = [−0.45 0.28]; *t*-tests).

Similar to the M_3_ model results, the prior choices weights (*β_prior_choices_*) were not significantly different from zero for three out of the four conditions (Fig. 6, second row, middle plot: vis→vis: *t*_19_ = 1.56, ***p* = 0.14**, *Cohen’s d* = 0.35, 95% *CI* = [−0.09 0.63]; vis→ves: *t*_19_ = 1.06, ***p* = 0.30**, *Cohen’s d* = 0.24, 95% *CI* = [−0.19 0.59]; ves→vis: *t*_19_ = 0.92, ***p* = 0.37**, *Cohen’s d* = 0.21, 95% *CI* = [−0.15 0.37]; *t*-tests), and were significantly positive for the ves→ves condition (*t*_17_ = 3.29, ***p* = 4.4∙10^−3^**, *Cohen’s d* = 0.77, 95% *CI* = [0.18 0.85]; *t*-test). Also, as expected, the current stimulus weight (*β_stimulus_*) was large and significantly positive in all four conditions (Fig. 6, second row, left plot: ves→ves: *t*_17_ = 10.34, ***p* = 9.5∙10^−9^**, *Cohen’s d* = 2.44, 95% *CI* = [2.98 4.50]; vis→vis: *t*_19_ = 8.20, ***p* = 1.2∙10^−7^**, *Cohen’s d* = 1.83, 95% *CI* = [6.58 11.09]; vis→ves: *t*_19_ = 9.45, ***p* = 1.3∙10^−8^**, *Cohen’s d* = 2.11, 95% *CI* = [2.77 4.35]; ves→vis: *t*_19_ = 5.13, ***p* = 5.9∙10^−5^**, *Cohen’s d* = 1.15, 95% *CI* = [5.02 11.92]; *t*-tests). The general bias weight (*β_0_*) was not significantly different from zero in any condition (Fig. 6, second row, right plot: ves→ves: *t*_17_ = 1.20, ***p* = 0.25**, *Cohen’s d* = 0.28, 95% *CI* = [−0.14 0.52]; vis→vis: *t*_19_ = 1.45, ***p* = 0.16**, *Cohen’s d* = 0.33, 95% *CI* = [−0.18 0.99]; vis→ves: *t*_19_ = 2.03, ***p* = 0.057**, *Cohen’s d* = 0.45, 95% *CI* = [−0.02 1.47]; and ves→vis: *t*_19_ = 1.41, ***p* = 0.18**, *Cohen’s d* = 0.32, 95% *CI* = [−0.19 0.99]; *t*-tests).

### Parameter recovery

The finding in the previous section that in the cross-sensory conditions the first stimulus prior (trial *t*-3) seemed to have a significant effect while the more recent stimulus priors (immediately preceding the test stimulus) did not, was somewhat surprising. To test that this was not a spurious result of the model fitting procedure, we performed further analysis using model simulations and parameter recovery. Specifically, we simulated performance for the task used in this study for each one of the 20 participants using their M_3E_ model fits. We also did this for one additional representative (fictitious) participant with predetermined model parameters (see Methods for further details). The simulated behavior (choices) was then analyzed in the same way as the data to extract (recover) the beta parameters. This procedure was repeated 100 times for each participant (resulting in 100 sets of 6 recovered betas per participant). We found that the recovered betas (median parameter values) were highly similar to and strongly correlated with the original betas (Suppl. Fig. 2, R^2^ ≥ 0.98).

In addition, we simulated the behavioral data also after reversing the order of the original prior stimulus parameters *β_prior_stimulus1,_ β_prior_stimulus2_*, and *β_prior_stimulus3_* (i.e., the original *β_prior_stimulus3_* was switched with *β_prior_stimulus1_*). We then repeated the parameter recovery process and successfully recovered the beta coefficients in this case too: the medians of the recovered betas were similar and strongly correlated with the original (reverse ordered) betas (Suppl. Fig. 3A, R^2^ ≥ 0.98). Finally, when comparing the original betas (not reversed) to the reversed recovered betas, the strong correlation in *β_prior_stimulus1_* and *β_prior_stimulus3_* was, as expected, eradicated (Suppl. Fig. 3B, R^2^ < 0.01). These results of the simulations and parameter recovery process indicate that the model fits for M_3E_ are reliable and unbiased.

## DISCUSSION

This study aimed to investigate whether a series of several short-duration stimuli might elicit cross-sensory adaptation. For this, we designed and ran an experiment that tested the effects of three short (1s) self-motion stimuli with biased headings (visual or vestibular) on subsequent (visual or vestibular) heading perception. We found significant adaptation of heading perception in both uni-sensory (visual and vestibular) as well as both cross-sensory (visual to vestibular, and vestibular to visual) conditions. This is the first demonstration of visual-vestibular (cross-sensory) adaptation for short-duration self-motion stimuli.

Cross-sensory adaptation of self-motion has been demonstrated before, but only for long-duration (15s) visual adapting stimuli (Cuturi & MacNeilage, 2014). Shorter duration visual stimuli (≤ 7.5s) did not lead to subsequent cross-sensory (vestibular) adaptation. In that study only visual→vestibular (i.e., not vestibular→visual) cross-sensory adaptation was tested. Crane (2013) also found no cross-sensory adaptation to short-duration (1.5s) self-motion stimuli (testing both visual→vestibular and vestibular→visual).

The difference between our results vs. these previous findings may stem from several factors. 1) In our experiment, three prior (‘adapting’) stimuli were presented; whereas in those experiments one adapting stimulus was presented. Accordingly, there may be a compound effect of several stimuli. This would not likely reflect a simple summation of stimulus duration, because when summed together, 3s of sensory information is still substantially less than 7.5s, which did not previously elicit cross-sensory adaptation (Cuturi & MacNeilage, 2014). Rather, it might reflect a higher-level compounding effect of sensory events. 2) In our study, both the adapting (prior) and the test stimuli were discriminated, whereas in both of the previous studies the participants were instructed to discriminate only the ‘test’ stimulus. Active discrimination may change a person’s experience, memory, and interpretation of stimuli (Gerrits & Schouten, 2004; Novak, Ritter, Vaughan, & Wiznitzer, 1990), which could lead to greater cross-sensory adaptation. 3) Although all three studies tested linear self-motion perception, the task in our study (heading discrimination, right or left of straight ahead) was slightly different vs. forward-backward discrimination from the other two studies. There is a possibility that heading discrimination is more susceptible to cross-modal adaptation.

Although we cannot dissociate the first explanation (a compounding effect of several discrete prior stimuli) from the second explanation (a boosting effect of active discrimination), both of these possible explanations indicate that the brain monitors and dynamically adapts to recent (discrete) sensory events in a supra-modal manner. Results from the model fits, that adaptation was primarily driven by prior stimuli (not choices), which might suggest that stimulus repetition (rather than active discrimination) is the driving factor. Future studies that test: i) only one, discriminated, prior or ii) three not-discriminated priors are needed to fully dissociate these factors. Also, future studies might investigate whether this result generalizes to other tasks.

In line with other visual and vestibular motion aftereffects (Crane, 2012; Cuturi & MacNeilage, 2014; Gordon et al., 1995; Kohn, 2007; Thompson, P., & Burr, 2009), we found that the prior trials had a ‘repulsive’ influence on subsequent perception. Namely, observers demonstrated a higher propensity for choosing left after rightward biased priors (and vice versa). This was seen for both of the unisensory conditions and both of the cross-sensory conditions. This repulsive effect seems different (opposite in direction) from the attractive effect often described as ‘serial dependence’ (Feigin et al., 2021; Fischer & Whitney, 2014; Liberman et al., 2014, 2018; Taubert & Alais, 2016).

Converging evidence suggests that these opposing effects (repulsive and attractive) of prior sensory experience reflect the independent influences of prior stimuli and prior choices, respectively. Namely, perceptual decisions are both attracted to prior choices and repulsed from prior stimuli (Bosch, Fritsche, Ehinger, & de Lange, 2020; Feigin et al., 2021; Feigin, Shalom-Sperber, Zachor, & Zaidel, 2020; Fritsche, Mostert, & de Lange, 2017; Sadil, Cowell, & Huber, 2021). Fitting our data with a logistic regression model (designed to dissociate the separate effects of prior stimuli and prior choices) provided further evidence in support of this view. Specifically, we found that adaptation in all four conditions (both uni-sensory and cross-sensory) was significantly driven by the prior stimuli, and not by the prior choices. A significant effect of prior choices was seen in one condition (ves→ves), and it was indeed attractive (in the opposite direction to the prior stimulus effect).

The Bayesian model comparisons indicate that stimulus history should be taken into account when modeling perception. There was very strong evidence for this in both unisensory conditions, and substantial evidence in one cross-sensory condition (vis→ves) but not the other (ves→vis; although the prior stimulus effect was still significant in that condition). This suggests that cross-sensory effects are not symmetrical and that vestibular perception may be more affected by visual events (than vice versa) – in line with findings of quicker vestibular vs. visual adaptation to multisensory conflict (Zaidel et al., 2011). This may reflect the fact that the primary visual cortices in primates are predominantly visual, with (less-dominant) non-visual signals joining higher up in the visual hierarchy (Van Essen, Anderson, & Felleman, 1992; Van Essen & Maunsell, 1983), whereas vestibular cortical regions are inherently multisensory from the outset (Brandt, 2003; Guldin & Grüsser, 1998). Overall, these results indicate that to model how the brain dynamically forms percepts we need to take into account that it is continuously affected by the statistics of (cross-modal) stimulus history.

Cross-modal aftereffects have been found also in other sensory modalities: visual motion stimuli cause cross-modal aftereffects in auditory perception (Ehrenstein & Reinhardt-Rutland, 1996; Kitagawa & Ichihara, 2002), and auditory motion can elicit a visual motion aftereffect (Berger & Ehrsson, 2016). Also, motion aftereffects transfer cross-modally between vision and touch (Konkle, Wang, Hayward, & Moore, 2009). In addition, the visual tilt aftereffect transfers to touch (Krystallidou & Thompson, 2016). This latter study further found that the visual-tactile tilt aftereffect occurred in spatiotopic (gravitational) coordinates, suggesting adaptation of high-level (perceptual) representations. Interestingly, cross-modal (audio-visual) effects extend even to emotional processing – sounds, like laughter, adapt the visual perception of emotional faces (Wang et al., 2017).

This growing literature on cross-modal aftereffects has important implications for understanding adaption in the brain. Taken together, it suggests that the brain exhibits high-level functional (perceptual) adaptation, beyond low-level (sensory) adaptation. Namely, cross-modal aftereffects likely reflect neuronal adaptation in cortical areas that are more process-dependent (i.e., functionally oriented) than modality-dependent (Konkle & Moore, 2009). This is also in line with high-level visual (unisensory) adaptation to global scene properties (Greene & Oliva, 2010). Our results are in line with this notion, and extend it by showing that discrete self-motion events (that may be too short to elicit sustained vection; Bubka, Bonato, & Palmisano, 2008; Seno, Ito, & Sunaga, 2010) still lead to cross-modal adaptation. They therefore suggest that the brain monitors statistics of perceptual events for specific functions (e.g. self-motion) supra-modally, biasing subsequent perception of that function, irrespective of its presented modality (e.g., visual or vestibular). This also means that unisensory adaptation (which is stronger than cross-modal) likely reflects a composite of function adaptation, and low-level (sensory) adaptation.

In the primate brain, there are multiple cortical areas that process both visual and vestibular cues of self-motion. These include the dorsal region of the medial superior temporal area (MSTd; F. Bremmer, Kubischik, Pekel, Lappe, & Hoffmann, 1999; Duffy, 1998; Gu, Watkins, Angelaki, & DeAngelis, 2006), the ventral intraparietal area (VIP; Frank Bremmer, Klam, Duhamel, Ben Hamed, & Graf, 2002; Chen, DeAngelis, & Angelaki, 2013), and the visual posterior sylvian area (VPS; Chen, DeAngelis, & Angelaki, 2011; Frank, Wirth, & Greenlee, 2016). An enigma of visual-vestibular responses in MSTd (and VIP) is the observation that alongside neurons with ‘congruent’ tuning (i.e., overlapping preferred directions, for visual and vestibular self-motion stimuli) many neurons have ‘opposite’ tuning (Chen et al., 2013; Gu, Angelaki, & DeAngelis, 2008). Thus, if cross-modal adaptation is attributed to overlapping multisensory responses in these neurons, e.g., via neuronal correlations (Schwartz, Hsu, & Dayan, 2007), then the effects on congruent and opposite neurons could perhaps cancel one another out (unless these different populations are selectively decoded for different purposes; Zhang et al., 2019). By contrast, visual self-motion responses in VPS are primarily tuned in the opposite direction (180° apart) vs. vestibular responses (Chen et al., 2011). Here, adaptation to visual cues might elicit cross-modal adaptation of vestibular responses in VPS. Further research of neuronal responses during cross-modal adaptation is required to better elucidate its neuronal basis.

In an exploratory analysis, we investigated the separate effects of the three prior stimuli using an expanded regression model. Not surprisingly, we found that uni-sensory adaptation was mainly affected by more recent sensory information (i.e., the two trials that preceded the test trial, *t*−1 and *t*−2). But, surprisingly, cross-sensory adaptation seemed to be mainly affected by the earliest of the three priors (*t*-3). We confirmed the validity of the model fits using simulations, and parameter recovery. However, because this was an exploratory analysis, we call for further research to confirm and to better understand it. We speculate that this finding may be due to increased salience of the stimulus at *t*-3 in the cross-sensory conditions. In the cross-sensory conditions, priors and test trials were of different sensory modalities. Due to this design, the first prior of a new batch (*t*-3 in relation to the test of that batch) came after a test trial of a different sensory modality (e.g., for ves→vis, the first vestibular prior of a batch came after a visual test stimulus). Since stimulus salience depends on novelty (among other factors; Downar, Crawley, Mikulis, & Davis, 2002; McDermott, Malkoc, Mulligan, & Webster, 2010; Sidlauskaite et al., 2014), it is possible that the novelty of the sensory modality made this prior more salient such that it had a greater effect on the test trial than the two following priors (*t*-2 and *t*-1).

In summary, we found here uni-sensory and cross-sensory (visual-vestibular) adaptation to several, discrete, short-duration (1s) self-motion stimuli. Adaptation was driven by the prior stimuli, not choices. These results suggest that the brain continuously monitors and adapts to the statistics of high-level percepts (supra-modally), leading to functional adaptation.

## AUTHOR CONTRIBUTIONS

S.S.S. analyzed the data and wrote the manuscript. A.C. contributed to writing the manuscript and A.Z. designed the study and wrote the manuscript.

## ACKNOWLEDGMENTS

We would like to thank Elad Goldberg and Orly Halperin for collecting the data and performing primary analyses. We would also like to thank Avraham Elkara for software development, David Swissa for mechanical and machinery development, and Tamar Harpaz for management assistance. This work was supported by a grant from The Israeli Centers of Research Excellence (I-CORE, Center No. 51/11) to A.Z. and by the ISF-NSFC joint research program (grant No. 3318/20) to A.Z. and A.C.

## SUPPLEMENTARY FIGURES

**Supplementary Figure 1.**
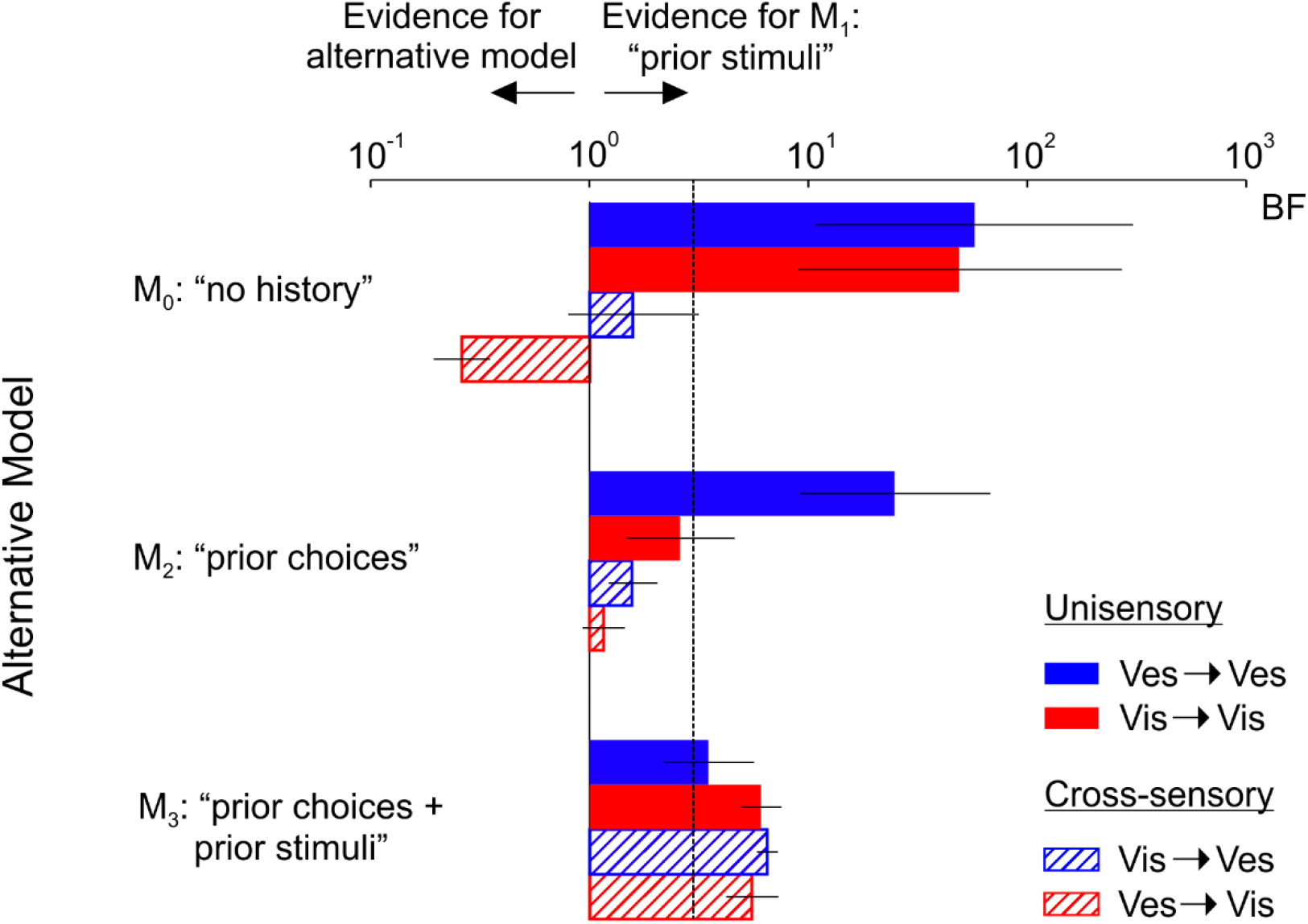
BIC Bayesian model comparisons. Related to Figure 5. All conventions are the same as in Figure 5. However, the Bayesian information criterion (BIC) was used instead of the Akaike information criterion (AIC).

**Supplementary Figure 2.**
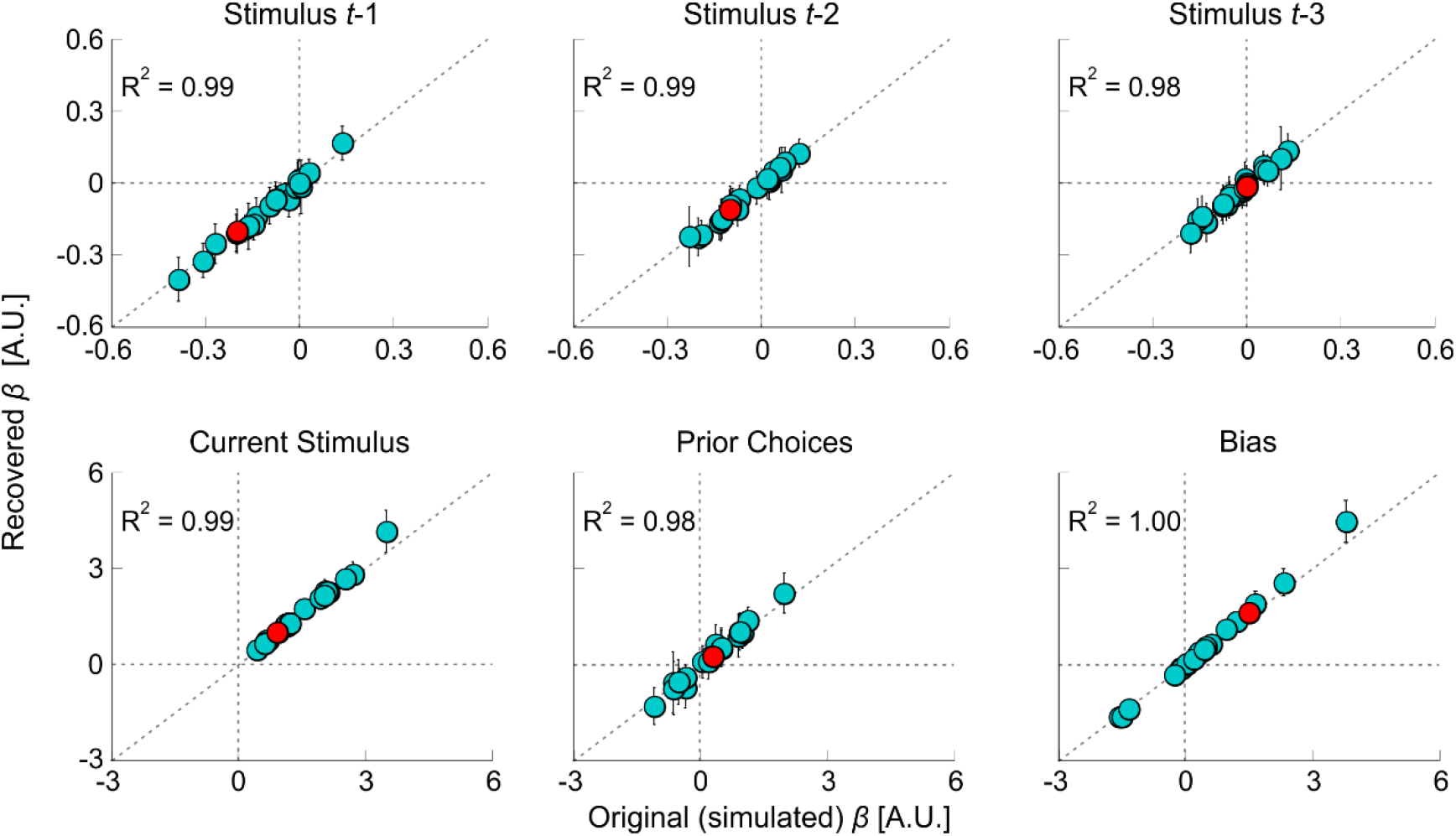
Parameter recovery. Related to Figure 6. The beta parameters for each participant (‘original’ betas, extracted from M_3E_) were used to simulate a new set of responses, following the same protocol as the experiment. ‘Recovered’ betas were then extracted from the simulated data. This was performed 100 times per participant. Each subplot (per parameter) presents the recovered betas (median ± MAD, median absolute difference) vs. the original betas. The cyan circles represent individual participants. The red circles represent simulated data for one fictitious participant. R^2^ values reflect the proportion of variance of the recovered betas’ medians that is explained by the original betas. Diagonal lines mark the line of equality (y = x).

**Supplementary Figure 3.**
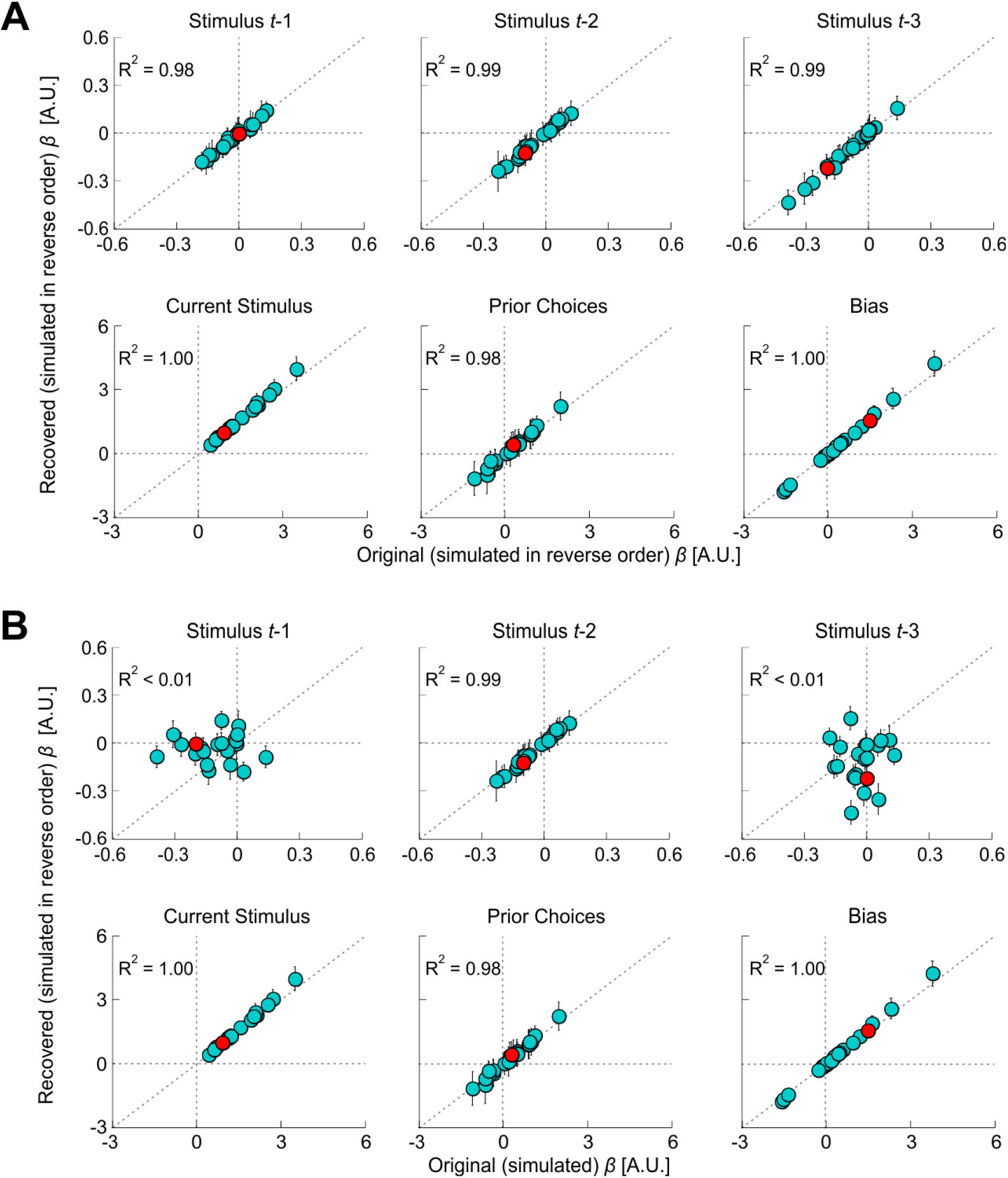
Parameter recovery after simulating with reversed order of prior stimuli. Related to Figure 6 and Supplementary Figure 2. All conventions are the same as Supplementary Figure 2. (A) The order of the prior stimulus betas (top row) was reversed, per participant, before simulation (i.e., *βprior_stimulus1* was switched with *βprior_stimulus3*), and then recovered. (B) The betas recovered from simulating in the reverse order (as in A) plotted vs. the original (not reversed) betas.

